# The effect of microbial metabolites from colonic protein fermentation on bacteria-induced cytokine production in dendritic cells

**DOI:** 10.1101/2024.07.17.603665

**Authors:** Zhuqing Xie, Danny Blichfeldt Eriksen, Peter Riber Johnsen, Dennis Sandris Nielsen, Hanne Frøkiær

## Abstract

**Scope:** Compared to the well-defined immune-modulating effect of butyrate, the knowledge of other metabolites from colonic protein fermentation is limited.

**Methods and results:** The effect of protein-derived metabolites (valerate, branched-chain fatty acids, ammonium, phenol, p-Cresol, indole, and H_2_S) on cytokine production in murine bone marrow-derived dendritic cells (BMDCs) stimulated with LPS, *Lactobacillus acidophilus* NCFM, or *Staphylococcus aureus* USA300 was investigated. The metabolites modulated the cytokine profile differently and depended on the specific microbial stimulus with short-chain fatty acids (SCFAs) exhibiting the strongest effects and no toxicity. After short-term treatment, SCFAs affected the cytokine profile similar to but weaker than butyrate, reflected by inhibition of IL-12p70 and IL-10 but enhanced IL-23 (LPS and *S. aureus* USA300) and IL-1β production. Compared to valerate, butyrate exhibited a stronger and more prompt effect on cytokine gene expression without influencing reactive oxygen species (ROS) formation. Oppositely, long-term treatment with the two SCFAs resulted in similar anti-inflammatory effects, i.e. abrogation of LPS-induced IL-12 and enhancement of IL-10 and the expression of aryl hydrocarbon receptor (*Ahr*) and LPS-stimulated dual specificity phosphatase 1 (*Dusp1*).

**Conclusion:** Our data reveals immune-modulating effects of various protein fermentation metabolites, and valerate in specific holds activities resembling but not identical to butyrate.

## 1. Introduction

Protein is an indispensable nutrient being essential for cellular functions, connective tissue, and muscle synthesis.^[1]^ However, the average protein consumption in most western countries exceeds the recommended intake (0.8–1.0 g/kg of body weight per day).^[2]^ Despite most protein being broken down and absorbed in the upper gastrointestinal tract, a certain amount (∼3-18 g/day)^[3, 4]^ of protein from diet and endogenous cells enters the colon and is fermented into various metabolites by the human gut microbes. In contrast to the short-chain fatty acids (SCFAs) production of fiber fermentation, protein fermentation in the distal colonic lumen contributes to not only SCFAs, but also results in branch-chain fatty acids (BCFAs), ammonia, hydrogen sulfide (H_2_S), p-Cresol, and phenolic and indolic compounds.^[5]^ These metabolites influence the human host in different ways. For instance, it was shown that ammonium^[6, 7]^ and phenolic compounds^[8–10]^ are detrimental, whereas indolic compounds from tryptophan fermentation can attenuate inflammation through activation of aryl hydrocarbon receptor (AHR) signaling.^[11, 12]^ The exact influence of H_2_S is unclear as conflicting results were reported.^[13, 14]^ Besides, the BCFAs isobutyrate, 2-methylbutyrate, and isovalerate are produced from the branched-chain amino acids valine, leucine, and isoleucine, respectively^[15]^ and have not been associated with negative influences. As one of the SCFAs, microbial fermentation-produced valerate has been reported to improve gut integrity at 2 mM^[16]^ and modulate lymphocytes^[17]^ and CD8^+^ T cell responses^[18]^ efficiently.

Dendritic cells (DCs) are antigen-presenting cells, acting as the bridge between innate and adaptive immunity where the cytokines produced in response to microbial stimuli determine the type of Th cells being activated.^[19–22]^ It is well-established that SCFAs especially butyrate, regulate the microbial-induced immune response through multiple pathways including G-protein coupled receptors (GPRs) and histone deacetylase (HDACs) inhibition.^[23]^ Butyrate influences the differentiation and maturation of DCs and downregulates the IL-12 production while increasing the production of IL-23 in LPS-stimulated DCs.^[24]^ Furthermore, long-term treatment of DCs with butyrate inhibited the production of IL-12.^[25]^ Compared to the widespread attention to the immune-modulating effect of the SCFAs (butyrate, propionate, and acetate), the knowledge regarding the immune regulation by other gut microbial metabolites like protein fermentation-derived products is limited. In the present study, we compared the impact of various protein fermentation products including isobutyrate, 2-methylbutyrate, isovalerate, valerate, ammonia, phenol, p-Cresol, indole, and sulfides on cytokine production using different microbial stimuli including LPS, UV-inactivated *Lactobacillus acidophilus* NCFM, or *Staphylococcus aureus* USA300 in BMDCs. Due to the most prominent but varying effect of the four fatty acids, we studied their effect in more detail and compared the effects to the well-described metabolite butyrate. We believe our results provide valuable information for understanding the potential influence of metabolites coming from proteinaceous substrates and serve as references for further in-depth explorations.

## 2. Results

### 2.1 Metabolites of protein fermentation affect BMDC viability

We first assessed the effect of the compounds at concentrations of 1 or 5 mM on BMDC viability (**Figure 1**). None of the fatty acids (isobutyrate, 2-methylbutyrate, isovalerate, and valerate) affected the BMDC viability (**Figure 1a-b**). In contrast, the rest of the compounds impacted the viability to varying degrees. Only phenol and p-Cresol at 5 mM caused cytotoxicity in cells stimulated with LPS (**Figure 1d**), but at 1 mM these compounds reduced cell viability when added to the immature BMDCs (**Figure 1c**). Also, indole reduced the viability of immature BMDCs to some extent.

**Figure 1:**
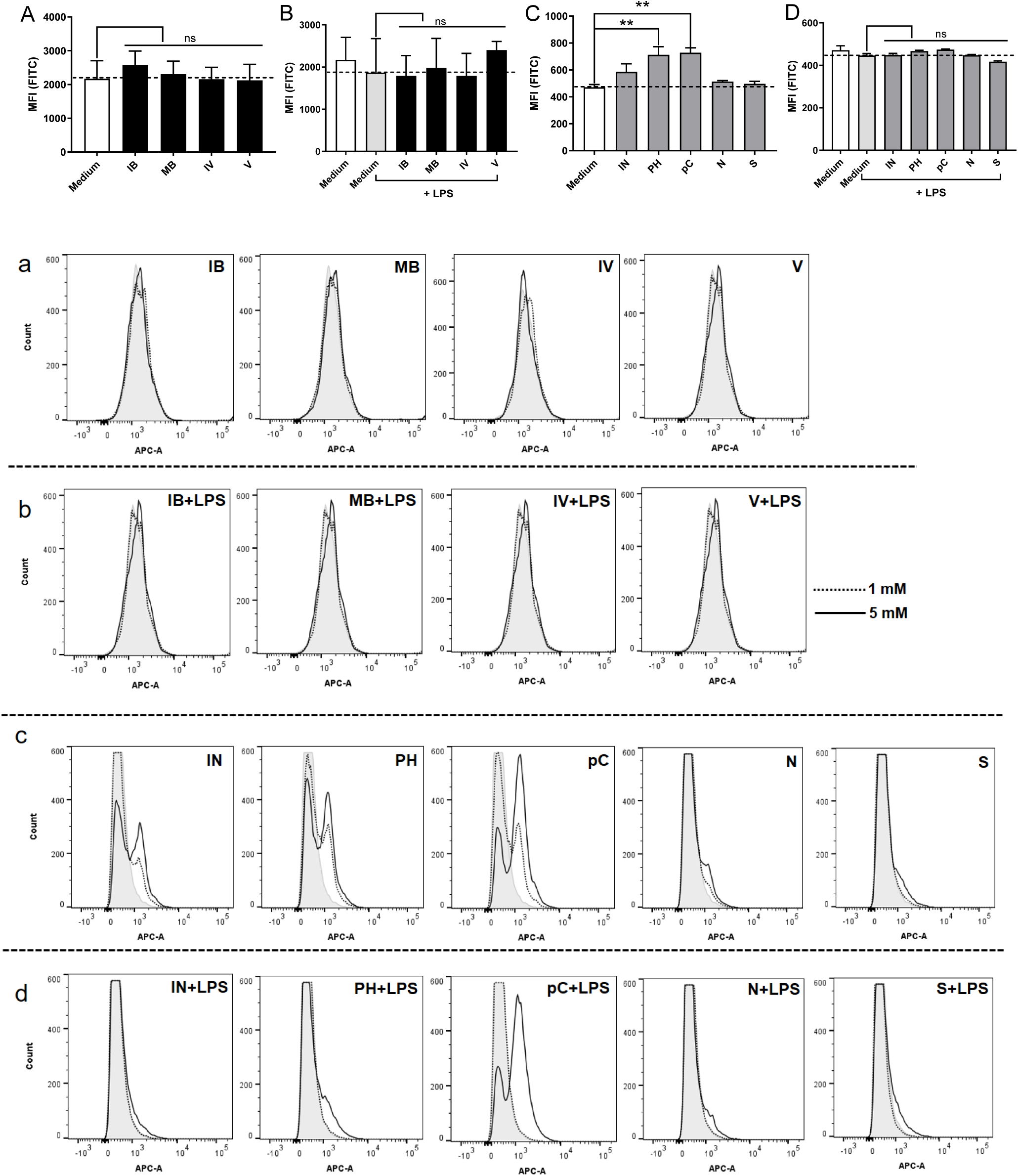
The influence of nine protein fermentation products on the viability of unstimulated or LPS-stimulated BMDCs. The compounds were added to BMDC 30 min prior to the stimulus LPS. After 20 hr incubation viability was assessed by addition of 7-AAD by flow cytometry. The median fluorescence index (MFI) of cells incubated with the various compounds (1 mM) is shown in **A** and **C** (unstimulated) and **B** and **D** (LPS stimulated). Representative flow cytometry histograms (1 mM or 5 mM) are shown in **a-d**. Differences in viability were assessed by one-way analysis of variance (ANOVA) followed by Dunnett (* p < 0.05, ** p < 0.01, *** p < 0.001, **** p < 0.0001, ns: not significant). IB, isobutyrate; MB, 2-methylbutyrate; IV, isovalerate; V, valerate; IN, indole; PH, phenol; pC, p-Cresol; N, ammonium chloride; S, sodium hydrosulfide.

### 2.2 Modulation of cytokine production by protein fermentation products depends on stimulatory agents (LPS, *L. acidophilus* NCFM, and *S. aureus* USA300)

When added to the BMDCs alone, none of the compounds induced the production of the cytokines investigated (results not shown). The effect of adding each of the compounds (1 mM) to the cells 30 min prior to microbial stimulation with LPS, *L. acidophilus* NCFM, or *S. aureus* USA300 on the production of IL-12, IL-1β, and IL-23 is shown in **Figure 2**, and the TNF-α, IL-6, and IL-10 production is shown in **Supplementary Figure 1**. Differences in the cytokine production were in particular evident for the IL-23, which was considerable after stimulation with LPS (>400 pg/ml) but low for *L. acidophilus* NCFM (<40 pg/ml), and of IL-1β (from ∼400 pg/ml after LPS or *S. aureus* USA300 stimulation to ∼40 pg/ml after *L. acidophilus* NCFM stimulation).

**Figure 2:**
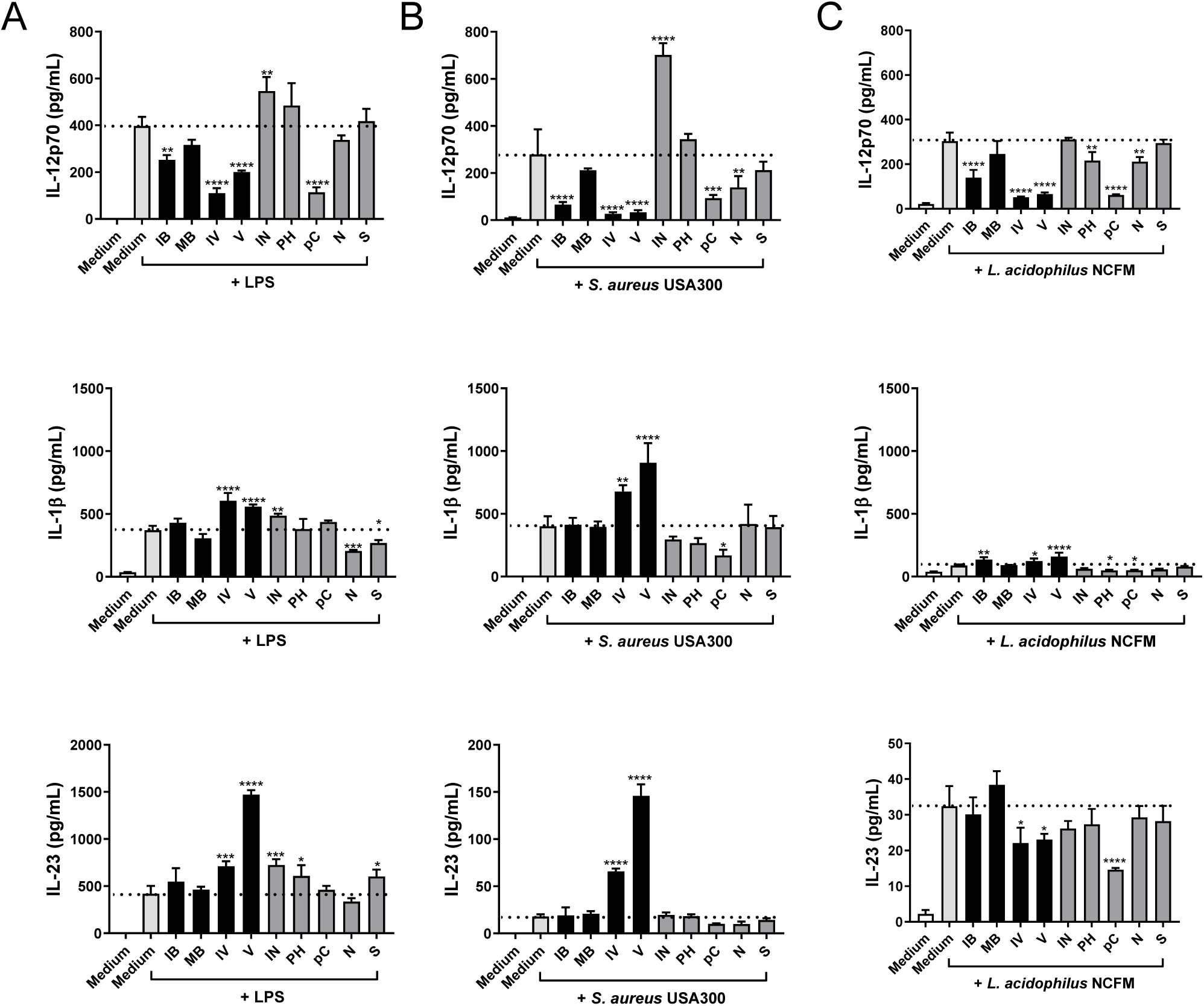
The effect of protein fermentation products on LPS **(A)**, *S. aureus* USA300 **(B)**, and *L. acidophilus* NCFM **(C)** induced IL-12p70, IL-1β, and IL-23 production. The compounds (1 mM) were added to BMDC 30 min prior to the bacterial stimuli. After 20 hr incubation supernatants were harvested and cytokine concentration was determined by ELISA after stimulation. Differences in the production of cytokine were assessed by one-way analysis of variance (ANOVA) followed by Dunnett (* p < 0.05, ** p < 0.01, *** p < 0.001, **** p < 0.0001). Y-axis coordinate range of IL-23 is different among stimulations. IB, isobutyrate; MB, 2-methylbutyrate; IV, isovalerate; V, valerate; IN, indole; PH, phenol; pC, p-Cresol; N, ammonium chloride; S, sodium hydrosulfide.

When stimulated with *L. acidophilus* NCFM (**panel C, Figure 2 and Supplementary Figure 1**), p-Cresol as the only metabolite reduced the production of all cytokines, isovalerate and valerate reduced IL-12, IL-23, and IL-10 and increased the production of IL-1β, and other metabolites affected the cytokine production but to a lesser extent. When stimulated with *S. aureus* USA300 (**panel B, Figure 2 and Supplementary Figure 1**), p-Cresol and ammonium chloride reduced the production of almost all cytokines, while indole enhanced the production of IL-12 and TNF-α, and fatty acids reduced the IL-12, IL-10, and IL-6 production while enhancing the IL-23 and IL-1β (only isovalerate and valerate). Together with LPS stimulation (**panel A, Figure 2 and Supplementary Figure 1**), all metabolites reduced IL-10 production, and some fatty acids further reduced IL-12 and TNF-α while enhancing the IL-23 and IL-1β. Overall, the SCFAs exhibited the most varying and distinct effect on the bacteria-induced cytokine production.

### 2.3 Compared to butyrate, some SCFAs show consistent and similar but weaker dose-dependent effects on bacteria-induced cytokines

We chose to assess the immunomodulating activity of the four SCFAs (isobutyrate, 2-methylbutyrate, isovalerate, and valerate) compared to butyrate in two different concentrations, 0.1 and 1 mM. BMDCs were pretreated for 30 min with the fatty acids prior to stimulation with LPS (**Figure 3**), *L. acidophilus* NCFM, or *S. aureus* USA300 (**Figure 4**), and the production of cytokines was determined. Taken together, each fatty acid showed a consistent dose-dependent but weaker influence on cytokine production compared to butyrate. Among the four fatty acids, valerate showed the strongest effect on modulating cytokine production.

**Figure 3:**
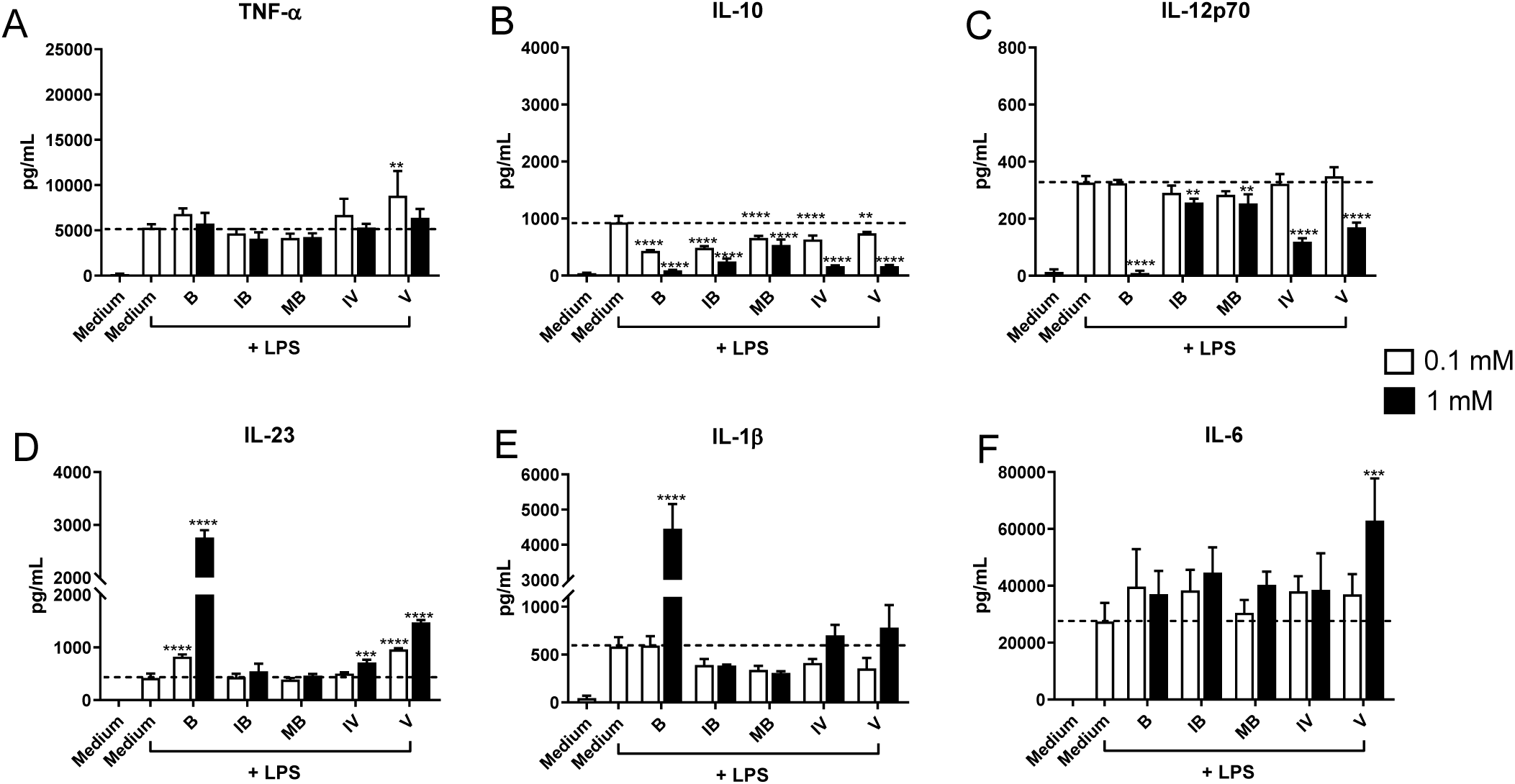
Immunomodulating dose-dependent effect of various fatty acids on LPS stimulated BMDC compared to butyrate. Different doses (1 mM vs 0.1 mM) of fatty acid were added to BMDC 30 min prior to stimulation with LPS (0.1 μg/mL). Cytokine production including TNF-α **(A)**, IL-10 **(B)**, IL-12p70 **(C)**, IL-23 **(D)**, IL-1β **(E)**, and IL-6 **(F)** after 20 hr was measured by ELISA. Differences in the production of cytokine were assessed by one-way analysis of variance (ANOVA) followed by Dunnett (* p < 0.05, ** p < 0.01, *** p < 0.001, **** p < 0.0001). Y-axis coordinate range of IL-23 and IL-1β is different among stimulations. B, butyrate; IB, isobutyrate; MB, 2-methylbutyrate; IV, isovalerate; V, valerate.

**Figure 4:**
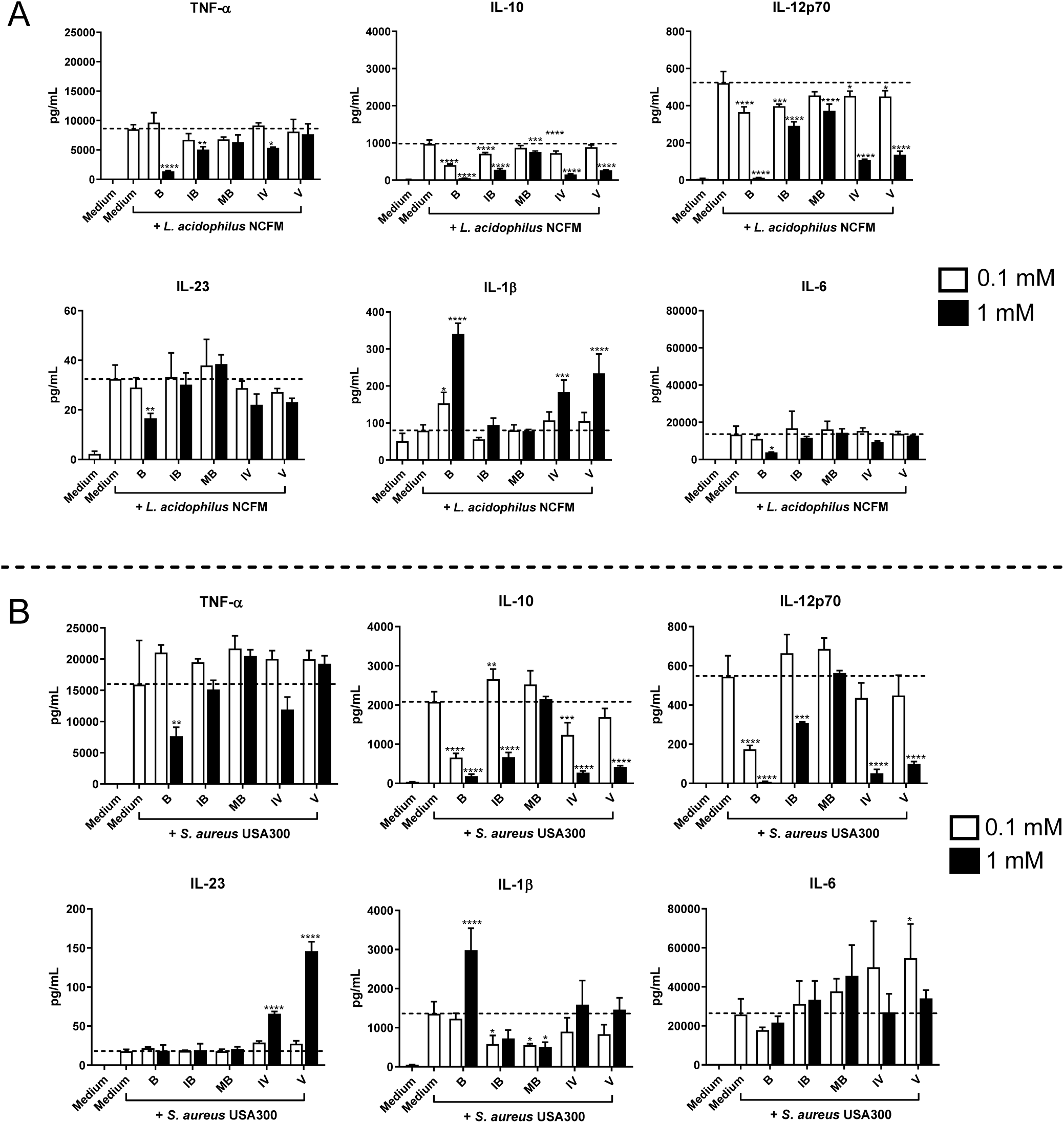
Immunomodulating dose-dependent effect of various fatty acids on bacteria stimulated BMDC compared to butyrate. Different doses (1 mM vs 0.1 mM) of fatty acid were added to BMDC 30 min prior to stimulation with *L. acidophilus* NCFM **(A)** and *S. aureus* USA300 **(B)**. Cytokine production including TNF-α, IL-10, IL-12p70, IL-23, IL-1β, and IL-6 after 20 hr was measured by ELISA. Differences in the production of cytokine were assessed by one-way analysis of variance (ANOVA) followed by Dunnett (* p < 0.05, ** p < 0.01, *** p < 0.001, **** p < 0.0001). Y-axis coordinate range of IL-23 and IL-1β is different among stimulations. B, butyrate; IB, isobutyrate; MB, 2-methylbutyrate; IV, isovalerate; V, valerate.

At 0.1 mM, each fatty acid inhibited the LPS-induced production of IL-10, whereas only butyrate and valerate promoted IL-23. All fatty acids at the high concentration of 1 mM reduced the LPS-induced IL-10 and IL-12 production, and butyrate, isovalerate, and valerate promoted the production of IL-23. Besides, butyrate and valerate at 1 mM led to high production of LPS-induced IL-1β and IL-6, respectively (**Figure 3**).

When stimulated with *L. acidophilus* NCFM (**Figure 4A**), most fatty acids at 0.1 mM significantly decreased IL-10 and IL-12, while only butyrate at 0.1 mM increased the production of IL-1β. At 1 mM, all fatty acids inhibited the production of IL-10 and IL-12, and some fatty acids decreased TNF-α but increased IL-1β. Butyrate was observed to have a downregulating effect on the production of TNF-α, IL-23, and IL-6 at 1 mM. Upon *S. aureus* USA300 stimulation (**Figure 4B**), butyrate at 0.1 mM decreased IL-10 and IL-12, and isobutyrate promoted IL-10 but decreased IL-1β. Valerate at 0.1 mM increased *S. aureus* USA300-induced IL-6 production. At 1 mM, only butyrate inhibited TNF-α, and most fatty acids but in particular isovalerate and valerate, decreased IL-10 and IL-12 but increased IL-23 production.

Together, butyrate and valerate showed the relatively strongest dose effect on modulating cytokine production among the fatty acids.

### 2.4 Valerate and butyrate have different effects on the ROS formation in BMDCs

The effect of adding butyrate or valerate to the cells 30 min prior to LPS stimulation on the ROS formation as determined by the oxidation of carboxy-H_2_DCFDA is shown in **Figure 5**. In non-stimulated cells, only valerate showed an influence on the ROS formation of BMDCs. After LPS stimulation, the ROS production increased as expected^[26]^ and was further increased by addition of valerate but not butyrate (**Figure 5 B, b**).

**Figure 5:**
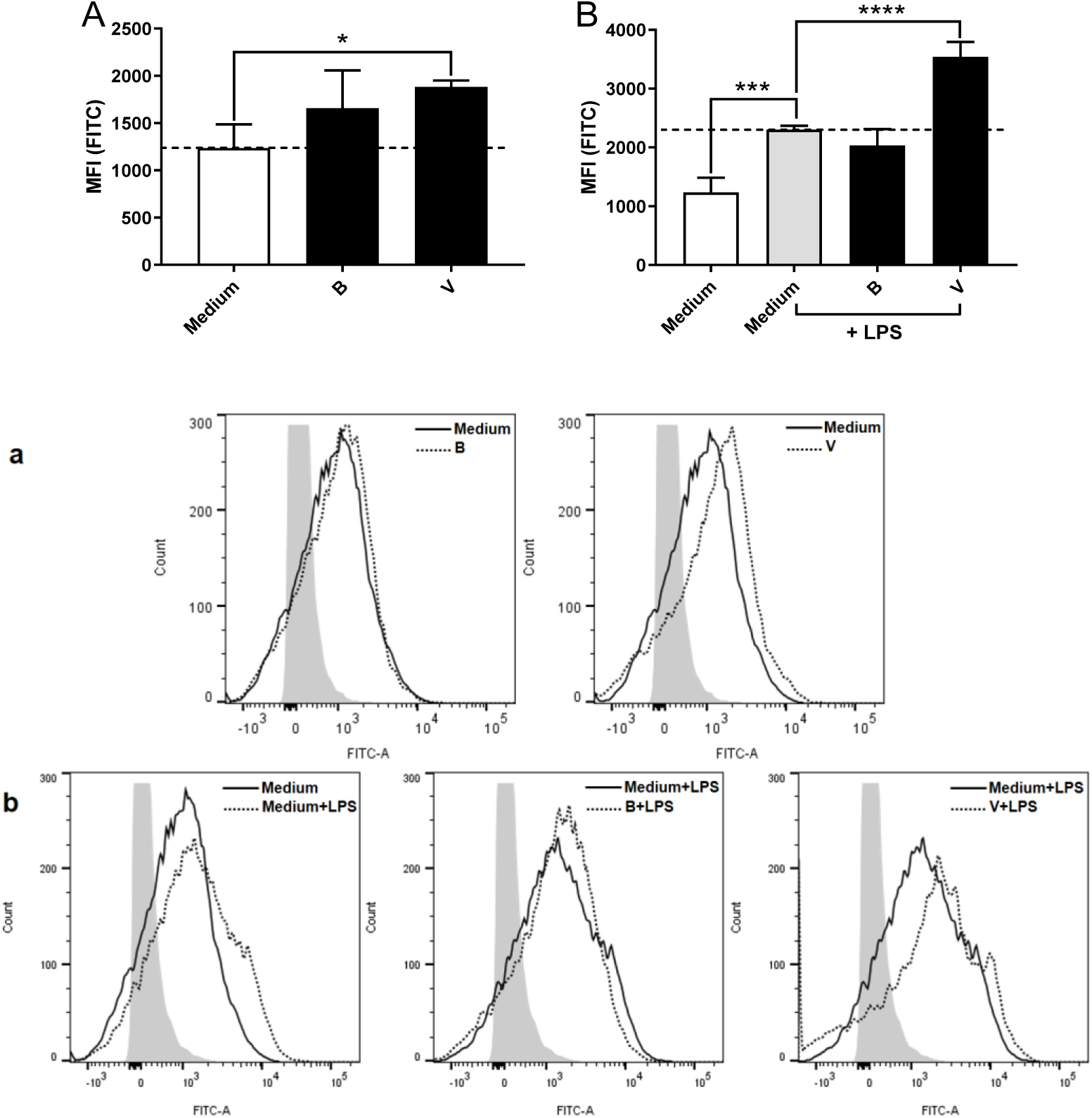
The influence of butyrate and valerate (1 mM) on ROS production with or without LPS stimulation (0.1 μg/mL). BMDCs were pretreated for 30 min with butyrate or valerate prior to stimulation with LPS, and the ROS formation was measured after 4 hr as the oxidation of carboxy-H_2_DCFDA by flow cytometry. The mean (n=3) fluorescence index (MFI) is shown in **A-B**, and representative flow cytometry results are shown in **a-b**. Differences in MFI were assessed by one-way analysis of variance (ANOVA) followed by Dunnett (* p < 0.05, ** p < 0.01, *** p < 0.001, **** p < 0.0001). B, butyrate; V, valerate.

### 2.5 Butyrate affects cytokine encoding gene expression more promptly than valerate

To assess how butyrate and valerate affected the expression kinetics of the cytokines IFN-β, IL-10, IL-12, and IL-23, BMDCs were pretreated with butyrate or valerate for 30 min prior to stimulation with LPS or *S. aureus* USA300 and expression of genes *Ifnβ*, *Il10*, *Il12a*, *Il12b*, and *Il23a* encoding for the respective cytokines, as well as the dual specificity phosphatase 1 (*Dusp1*), were measured after 2 (only LPS stimulation), 4, 6 and 8 hr, and the cytokines in the supernatant were quantified (**Figure 6**). Cells treated with LPS gave a more prompt response relative to *S. aureus* USA300 stimulation as reflected by the very early increase in *Ifnβ* and *Il12a* expression peaking after 2 hr compared to 6 hr or later after *S. aureus* USA300 stimulation. In contrast, the expression of *Il10* and *Il23a* peaked later; after 6 and 8 hr, respectively, and with no difference between the two stimuli. Addition of butyrate resulted in an almost complete abrogation of both LPS and *S. aureus* USA300-induced IFN-β, IL-12, and IL-10 production. In contrast, valerate only led to a partial reduction in IFN-β and IL-12 production, especially after LPS stimulation where the suppression was only modest. The different effects of butyrate and valerate on the LPS and *S. aureus* USA300-induced IFN-β and IL-12 and corresponding genes were especially evident at the early time points after stimulation. One key mechanism in the abrogation of the microbially stimulated IFN-β and IL-12 in BMDCs is increased *de novo* synthesis of *Dusp1* that efficiently dephosphorylates JNK and P38.^[27–29]^ Strikingly, the expression of *Dusp1* was promptly upregulated after butyrate but not after valerate treatment.

**Figure 6:**
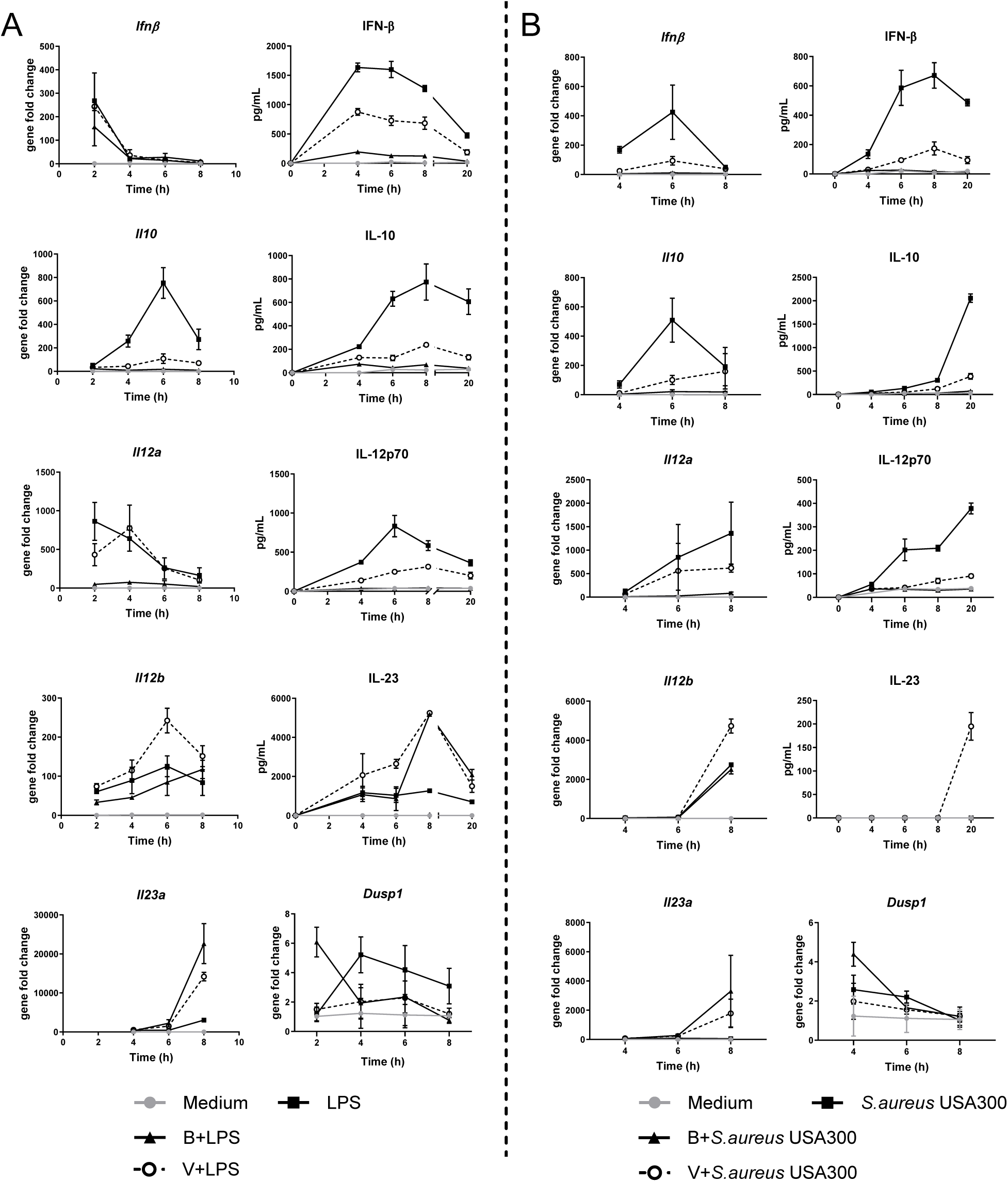
The influence of butyrate and valerate on the expression kinetics of the cytokines (IFN-β, IL-10, IL-12p70, and IL-23) and the encoding genes (*IfnB, Il10*, *Il12a, Il12b, Il23a,* and *Dusp1*) in BMDCs stimulated with LPS **(A)** or *S. aureus* USA300 **(B)**. BMDCs were pretreated for 30 min prior to stimulation with LPS or *S. aureus* USA300. *Actb* (Mm00607939_s1) was used as the reference gene to determine the ΔCt of each sample (ΔCt_target_ = Ct_target_ − Ct_reference_). The fold gene expression increase of the samples was analyzed by subtracting the ΔCt of the unstimulated cells (control) (ΔΔCt = ΔCt_target_ − ΔCt_control_) followed by the 2^(−(ΔΔCt)^ method. B, butyrate; V, valerate.

### 2.6 Long-term pretreatment of BMDCs with butyrate or valerate inhibits IL-12 and enhances IL-10 and TGF-β production

As cells may be conditioned by dietary fermentation metabolites long in advance to microbial stimulation, we aimed to assess the effect of adding butyrate or valerate two days before the fully differentiated DCs were harvested (pre-treatment) and compare this treatment with the effect of adding valerate or butyrate 30 min before stimulation with LPS (**Figure 7**). Expectedly, pre-treatment of the cells with either butyrate or valerate without any microbial stimuli did not give rise to the production of IL-12, IL-10, or IL-23. However, we also quantified TGF-β, which is constitutively expressed by immature DCs, and TGF-β raised to the level seen after stimulation with LPS (**Figure 7A**). When butyrate or valerate was added two days before stimulation with LPS, the induced IL-12 was almost completely abrogated, IL-10 was further increased, while the IL-23 and TGF-β production was unaffected (**Figure 7A**). Together, this indicates that both butyrate and valerate added during the late period of the DC development imprint a tolerogenic phenotype. When adding butyrate or valerate just before stimulation with LPS to cells pre-treated with butyrate or valerate, butyrate strongly suppressed the induced IL-10 and IL-23, while valerate only reduced IL-10 and only to the level induced by LPS (**Figure 7B**). Adding valerate to cells pre-treated with butyrate (**Supplementary Figure 2**) and stimulated with LPS resulted in cytokine production comparable with cells pretreated with valerate, thus further demonstrating that the effects of pre-treatment with the two SCFAs were comparable, while the short-term effects were markedly different.

**Figure 7:**
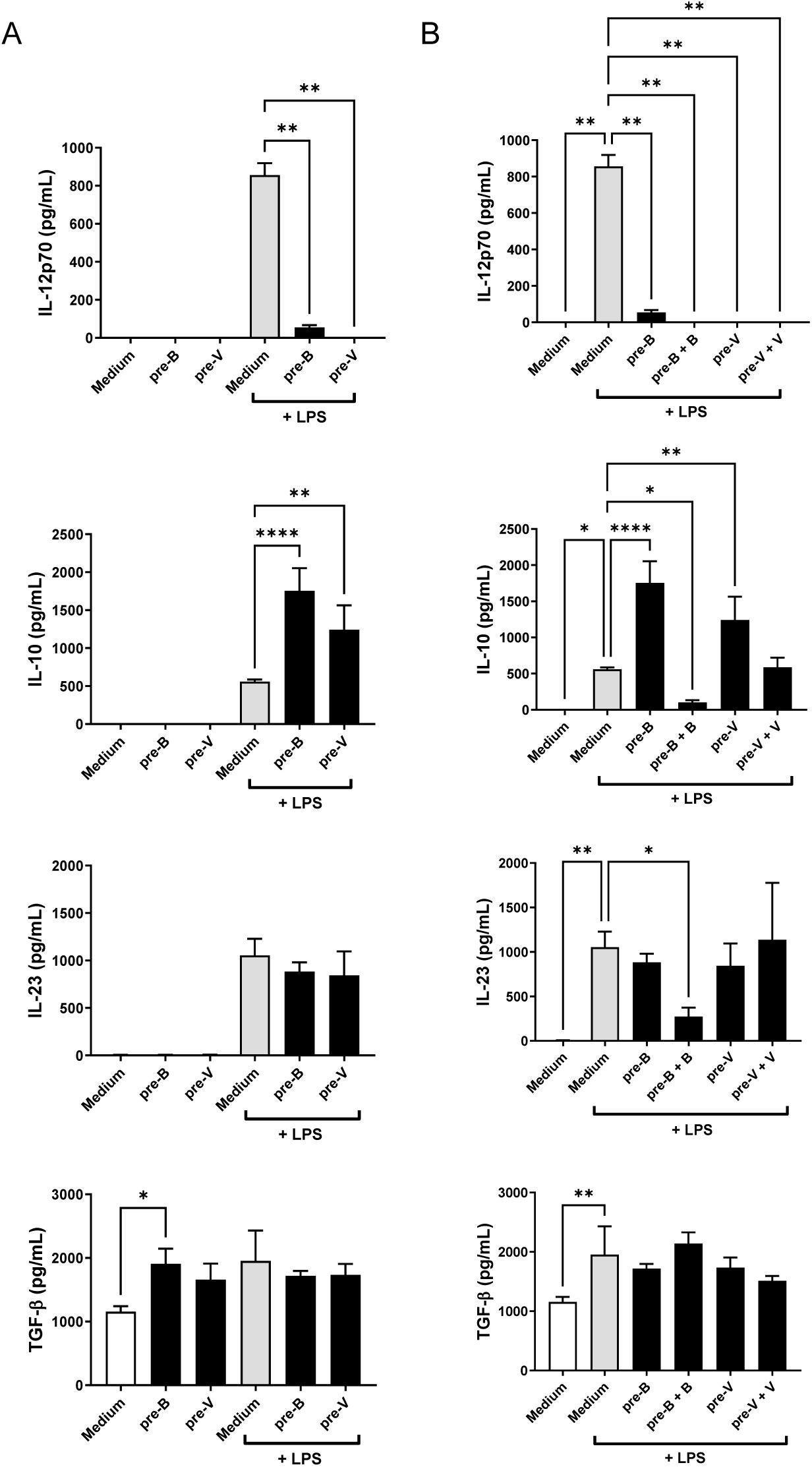
The production of cytokines in BMDCs pre-treated with butyrate (pre-B) or valerate (pre-V) for 2 days prior to harvest of cells and stimulation with LPS **(A)** and when further added butyrate (B) or valerate (V) 30 min before stimulation with LPS **(B)**. Cells were incubated for 20 hr and supernatants were harvested and analyzed for cytokine concentrations. Statistics: one-way analysis of variance (ANOVA) with multiple comparisons. **(A)**: Šídák’s multiple comparisons test. Asterisks show differences from media or LPS, respectively, with no added fatty acids. **(B)**: Dunnett’s multiple comparisons test. Asterisks show differences between LPS-stimulated cells with no added fatty acids. * p < 0.05, ** p < 0.01, *** p < 0.001, **** p < 0.0001.

A number of genes have previously been demonstrated to be involved in the regulation of microbial-induced cytokine production and function of dendritic cells, including *Ahr*,^[30]^ *Cyp1a1*,^[31]^ indoleamine 2,3dioxygenase 1 (*Ido1*),^[32]^ and *Dusp1.* AHR is indispensable for the induction of IL-10, which in turn inhibits the production of proinflammatory cytokines.^[33, 34]^ Long-term treatment with both butyrate and valerate enhanced the expression of *Ahr* in the immature BMDCs, while LPS reduced the level of *Ahr* expression (**Figure 8A**). The LPS-induced expression of IL-10 after long-term treatment (**Figure 7A**) corresponded with the increased *Ahr* expression. This pattern was also reflected in the expression of *Cyp1a1* upon butyrate treatment, although gene expression was weak 8 hr after LPS stimulation (**Figure 8B**). Furthermore, pre-treatment of the cells for two days led to an increase in the *Ido1* expression in non-stimulated cells but not in LPS-stimulated cells (**Figure 8C**). In contrast, the expression of *Dusp1* (**Figure 8D**) was upregulated in LPS-stimulated cells but not in unstimulated cells upon pre-treatment with the fatty acids.

**Figure 8:**
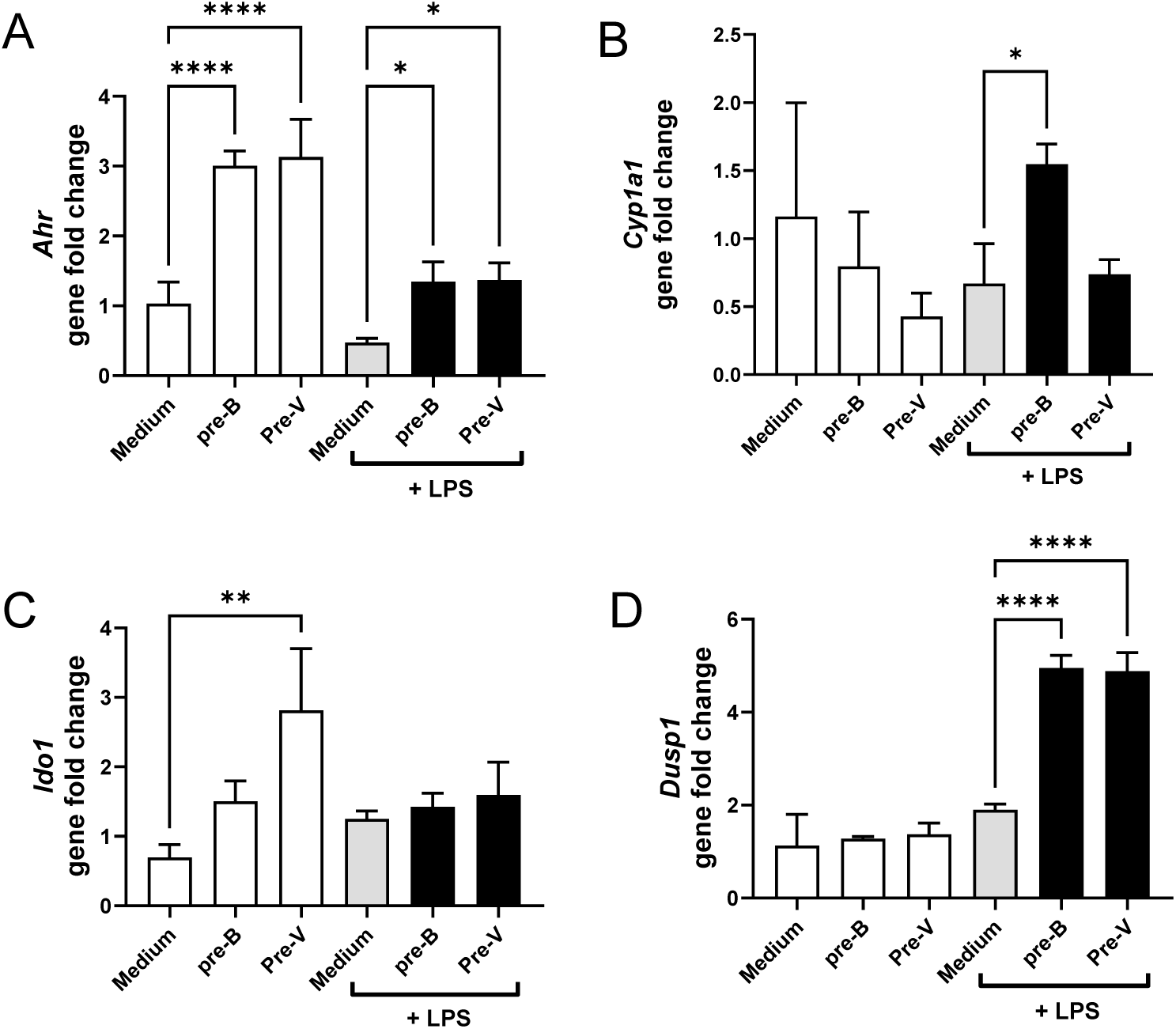
The expression of *Ahr* **(A)**, *Cyp1a1* **(B)**, *Ido1* **(C)**, and *Dusp1* **(D)** in BMDCs pre-treated with butyrate (pre-B) or valerate (pre-V) for 2 days prior to stimulation with LPS or media and incubated for 8 hr before cell harvest and isolation of mRNA. *Actb* (Mm00607939_s1) was used as the reference gene to determine the ΔCt of each sample (ΔCt_target_ = Ct_target_ − Ct_reference_). The fold gene expression increase of the samples was analyzed by subtracting the ΔCt of the unstimulated cells (control) (ΔΔCt = ΔCt_target_ − ΔCt_control_) followed by the 2^(−(ΔΔCt)^ method. Differences in gene fold change were assessed by one-way analysis of variance (ANOVA) with Šídák’s multiple comparisons. Asterisks show differences from media or LPS, respectively. * p < 0.05, ** p < 0.01, *** p < 0.001, **** p < 0.0001.

## 3. Discussion

A small but significant part of ingested proteins reaches the large colon undigested and may here become microbially fermented with the amino acids being converted into various metabolites by gut microbes. Compared to dietary fiber fermentation resulting in high production of SCFAs like butyrate which has barrier-enhancing and anti-inflammatory functions,^[35–37]^ the knowledge regarding other metabolites from protein fermentation is scarce with only the changes in epithelial integrity previously reported.^[4, 38, 39]^ Here, we examined the potential effect of nine of these metabolites (SCFAs/BCFAs, indole, phenol, p-Cresol, ammonium, and H_2_S) regarding their toxicity and their ability to modulate the induction of cytokine via various microbial stimuli (LPS, *L. acidophilus* NCFM or *S. aureus* USA300) in BMDCs. Of these compounds, the investigated SCFAs (isobutyrate, methylbutyrate, valerate, and isovalerate) exhibited low toxicity and the strongest immunomodulatory activity. We therefore investigated their effect on the BMDCs in more detail and found interesting differences in the modulating mechanisms.

All the compounds tested, except for SCFAs, affected the viability of immature BMDCs but to a varying degree. p-Cresol in particular exhibited high toxicity, both when added to immature and to LPS-stimulated BMDCs, and this correlated with a reduced cytokine production for all cytokines upon p-Cresol addition, thus indicating that the toxic effect influenced the cytokine production. The harmful effects of p-Cresol^[40]^ and phenol^[8]^ were also reported in previous studies on colonic epithelial cells where a concentration of 1 mM of phenolic compounds strongly impacted the viability. When assessing the effect on the cell viability in LPS-stimulated (mature) DCs, the toxic effect of p-Cresol was still high indicating a continuous killing of cells during or after maturation. This contrasts with the effect of many of the other compounds, which showed low proportions of dead cells after the addition of LPS. This may indicate that maturation by LPS, which induces increased endocytotic activity,^[41]^ could facilitate the clearing of dead cells through endocytosis. The effect on mature cells by p-Cresol indicates that this compound prevents (fully or partly) maturation by LPS. We did not, however, investigate this further. Indole-treated cells revealed only minor toxicity but nevertheless significantly reduced the microbially induced production of IL-10 and enhanced the *S. aureus* USA300 and LPS-induced IL-12. Indole and its derivatives are derived from the metabolism of tryptophan by gut microorganisms.^[42]^ Here we only tested indole and not the various derivatives which may exert different strengths and effects.

Our data do not allow us to distinguish between effects caused by cell toxicity and effects caused by changes in cell signaling. Accordingly, we chose to focus the study on SCFAs/BCFAs, which demonstrated no or very low toxicity. It is well-established that different microbial stimulations cause differences in the resulting cytokine production by DCs due to activation of distinct signaling pathways in the cell,^[22, 43]^ but it is not clear to which extent the effects of the metabolites depend on the microbial signal. In contrast to the other metabolites investigated, the tested SCFAs exhibited quite varying effects in the differently stimulated cells. Especially for the production of IL-23, TNF-α, and IL-6, we observed marked differences depending on the microbial stimuli. We have previously shown similar dependences on the microbial signal for the carbohydrates mannan and β-glucan^[44]^ as well as for stilbenoids.^[45]^ Hence, the type of stimulation should be considered when assessing the immunomodulatory effect of SCFAs *in vitro*.

Comparing two different doses of SCFAs with the effect of the same doses of butyrate, revealed that independently of the applied microbial stimuli, butyrate stood out as the SCFA with the strongest impact on the cytokine production. Isobutyrate, isovalerate, and especially valerate also exhibited potent immunomodulation while methylbutyrate was poor, indicating the importance of flexibility in the aliphatic chain close to the α-carbon for the activity of the SCFAs. One difference between the immunomodulating effects of the SCFAs seems to depend on the applied microbial stimuli. The IL-23 production increased dramatically in butyrate-treated cells after LPS stimulation, while after *S. aureus* USA300, but not *L. acidophilus* NCFM stimulation, valerate (and isovalerate) treatment revealed potent IL-23 increasing capacity. The enhanced IL-23 production seen after addition of butyrate to LPS-stimulated cells has been reported previously,^[24]^ but the lack of enhanced IL-23 production after stimulation with whole bacteria has to our knowledge not been described before. An important difference between stimulation with LPS and the intact gram-positive bacteria is that whereas LPS can induce an intracellular signaling cascade promptly by ligating to TLR4,^[46]^ the gram-positive bacteria must be taken up and degraded endosomally in order to release TLR-activating ligands,^[20]^ a process that results in a markedly delayed response as seen by the late upregulation of cytokine gene transcription. Thus, depending on whether the metabolites affect the cells promptly through receptor binding and subsequent intracellular signaling modification or, more slowly, through epigenetic modification, this may affect the cytokine production differently. To this end, butyrate, but not valerate, showed prompt upregulation of *Dusp1* expression encoding the dual specificity phosphatase 1, which deactivates the MAP kinases p38 and JNK.^[47]^ We have shown that induction of the production of IL12 and IFN-β depends on p38, JNK, or both kinases.^[22]^ Thus, butyrate may through the early *Dusp1* upregulation inhibit the early transcription of IL-12 and IFN-β encoding genes. As the induction of IFN-β by *S. aureus* USA300 is slower, the early upregulation of *Dusp1* by butyrate may therefore have less impact on the response.

Endosomal ROS production is a prerequisite for the degradation of endocytosed proteins and phagocytosed intact bacteria like *S. aureus* USA300 to produce cytokines.^[48, 49]^ The increased ROS formation induced by valerate but not by butyrate (with or without LPS stimulation) may be a factor involved in the divergent cytokine responses seen for the two SCFAs. However, as we did not determine the source of ROS we cannot know whether the increased ROS induced by valerate is induced through the activation of NADPH oxidase or by an effect of valerate on the mitochondrial ROS production as fatty acids have been demonstrated to have the capacity to affect both depending on the fatty acid as well as the cell type.^[50]^ Butyrate reportedly causes intracellular potassium ion outflow and hyperpolarization and calcium ion inflow, leading to activation of the NLRP3 inflammasome upon binding to GPR43 or GPR109a,^[51]^ which leads to increased IL-1β production. With no known receptor for valerate, an alternative explanation for the increased IL-1β could be the increased ROS production by valerate, as ROS is a known inducer of the NRLP3 inflammasome.^[52]^ This, however, is purely speculative and needs further investigation.

In contrast to the short-term effect assessed when adding the metabolites 30 min prior to microbial stimulation, long-term (2 days) treatment of the cells revealed comparable effects of butyrate and valerate on the cytokine production and was different from short-term treatment, resulted in an upregulation of IL-10 response in the LPS-stimulated BMDCs. Together, these differences between short- and long-term treatment may indicate a difference in whether there is an involvement of receptor-ligand interaction, such as the well-described interaction between GPR109a and butyrate in DCs,^[53]^ or not. Long-term treatment with both butyrate and valerate led to an increase in TGF-β corresponding to the increase seen after LPS stimulation (around 50% increase) (without *per se* inducing the production of other cytokines) and increased expression of *Ahr* and *Ido1* expression. The transcription factor AHR is involved in the induction of IL-10 upon LPS stimulation downregulating the production of pro-inflammatory cytokines^[54]^ and it further mediates the expression *of Ido1*.^[55]^ It has previously been demonstrated that in a microenvironment dominated by immunoregulatory TGF-β, IDO-1 becomes phosphorylated, which in turn leads to further upregulation of the expression of *Ido1* and *Tgfb1* genes,^[56]^ ultimately establishing a long-term immunoregulatory DC phenotype. In contrast to *Ido1* and *Ahr*, the expression of *Dusp1* was increased only upon LPS stimulation, underscoring its effect in newly stimulated cells. This is confirmed by the changed effect of LPS stimulation after both butyrate and valerate long-term treatment leading to increased IL-10 as opposed to short-term treatment with butyrate showing the most prominent effect leading to a diminished IL-10 production.

## 4. Conclusion

In summary, we have investigated the immunomodulatory role of a series of metabolites arising after protein fermentation by the gut microbiota in DCs. The SCFAs stood out as having low toxicity and high capacity to change the cytokine production of microbial-stimulated DCs with the unbranched SCFA having the strongest effect. The type of microbial stimulation (LPS, *L. acidophilus* NCFM, or *S. aureus* USA300) influenced the effect of the SCFA. The unbranched SCFA valerate had the strongest influence but compared to butyrate, the effect was weaker after short-term treatment. Long-term treatment with both butyrate and valerate led to a tolerogenic phenotype of DCs, showing both similarities and differences between the mechanisms involved in the way butyrate and valerate affect the cytokine production in BMDCs.

## 5. Materials and methods

### 5.1 Protein/amino acid-derived metabolites preparation

Ten compounds (indole, phenol, p-Cresol, ammonium chloride, sodium hydrosulfide, iso-butyrate, 2-methylbutyrate, iso-valerate, valerate, and sodium butyrate) were purchased from Sigma-Aldrich (St. Louis, MO, USA). The stock preparation was in sterile Milli-Q water except indole which was dissolved in ethanol. Final concentrations of ethanol never exceeded 0.1% on the cells. To exclude the impact of pH, NaOH at 50 mM was used in stock preparation when needed. Stock solutions were further diluted with cell culture medium at desired concentrations for treatments.

### 5.2 Bacterial strains preparation

Two gram-positive bacteria *L. acidophilus* NCFM (Danisco, Copenhagen, Denmark) and clinical methicillin-resistant *S. aureus* strain USA300 used in this study were prepared according to the previous report.^[44, 45]^ *L. acidophilus* NCFM grew anaerobically in de Man Rogosa Sharp broth (Merck, Darmstadt, Germany), whereas USA300 was streaked on tryptic soy agar and grown in tryptic soy broth at 37 ℃. Bacteria were washed twice in sterile PBS and seeded in Petri dishes for CFU determination. UV pulsation (>90 seconds; 6 s/pulse with a monochromatic wavelength of 254 nm; CL-1000 crosslinker; UVP, Cambridge, United Kingdom) was used to kill the bacteria, and the viability was verified afterward.

### 5.3 BMDCs generation, stimulations, and treatments

BMDCs were generated from C57BL/6NTac mice (Taconic, Lille Skensved, Denmark) as described previously.^[57, 58]^ Briefly, cells were isolated from the tibia and femur by flushing the bones with cold PBS, after which cells were seeded at 3 ∗ 10^5^ cells/mL in RPMI 1640 medium containing 10 % (v/v) heat-inactivated FBS, L-glutamine (4 mM), penicillin (100 U/ mL), streptomycin (100 U/mL), and 2-mercaptoethanol (50 μM). On days 3 and 6, culture medium containing 15 ng/ml granulocyte-macrophage colony stimulating factors (GM-CSF) was added to Petri dishes for DC differentiation. Non-adherent BMDCs were harvested on day 8 and diluted into 2 ∗ 10^6^ cells/mL, after which prepared compounds at desired concentrations were added for pre-incubation for 30 min at 37 ℃ in 5% CO^2^. LPS (0.1 μg/mL, *E. coli* O26:B6, Sigma Aldrich, St Louis, MO, USA), UV-treated *L. acidophilus* NCFM (MOI 1), or *S. aureus* USA300 (MOI 10) was added to simulate inflammation afterward. For the long-term treatment, BMDCs were pre-treated with butyrate (pre-B) or valerate (pre-V) for 2 days prior to stimulation with LPS.

### 5.4 BMDCs viability assay with flow cytometry

The effects of protein fermentation compounds on cell viability were assessed by calculating the intensity of the cellular fluorescence produced from 7-Aminoactinomycin D (7-AAD) using a BD FACS Diva flow cytometer (BD Biosciences, Franklin Lakes, NJ, USA). In brief, cells were washed with cold PBS after 20 hr of incubation and stained with 7-AAD for 1-3 hr, after which the dead cells were measured on flow cytometer with the calculation of the mean fluorescent intensity (MFI) using the FlowJo™ software (version 10.6.2, BD Life Sciences, Ashland, OR, USA). All events counted at 20,000/sample for the analysis.

### 5.5 Cytokine secretion assay

Cytokine production assay was performed in collected supernatants after 20 hr of incubation. Murine cytokines including TNF-α, IL-6, IL-10, IL-12p70, IL-23, and IL-1β were detected using duoset ELISA (R&D systems, Minneapolis, MN, USA) according to the manufacturer’s manual.

### 5.6 Determination of intracellular ROS formation

ROS formation was determined by measuring the intensity of the cellular fluorescence produced from the oxidation of the internalized carboxy-H_2_DCFDA on a BD FACS Diva flow cytometer (BD Biosciences, Franklin Lakes, NJ, USA). Briefly, murine BMDCs at 2 × 10^6^ cells/mL were stained with 5 μM carboxy-H_2_DCFDA (Invitrogen™, Carlsbad, CA, USA) and pre-incubated with prepared fatty acids (1 mM) at 37℃ and 5% CO_2_ for 30 min following the addition of 0.1 μg/mL LPS. BMDCs treated with fatty acids or LPS only as well as unstained BMDCs were listed as controls. After another 4 hr of incubation, the ROS formation was determined immediately, and the differences among fatty acids were assessed by calculating the mean MFI of treated BMDCs using the FlowJo™ software (version 10.6.2, BD Life Sciences, Ashland, OR, USA) with all events counted at 20,000/sample.

### 5.7 RNA extraction, cDNA synthesis, and quantitative real-time qPCR

RNA from BMDCs after treatments at different time points (2-8 hr) was extracted using the MagMAX-96 Total RNA Isolation Kit (Applied Biosystems, Foster City, CA, USA) according to the manufacturer’s instructions. cDNA was converted from RNA using the Applied Biosystems™ High-Capacity cDNA Reverse Transcription Kit. Actb (Mm00607939_s1) was used as the reference gene to determine the ΔCt of each sample (ΔCt_target_ = Ct_target_ − Ct_reference_). The fold gene expression of the samples was analyzed by subtracting the ΔCt of the unstimulated cells (ΔΔCt = ΔCt_target_ − ΔCt_control_) followed by the 2^(−(ΔΔCt)^ method.

### 5.8 Statistics

All the experiments in this study were carried out with at least three biological replicates including three technical repeats each time. Data are shown as means ± SD and analyzed using GraphPad Prism software (version 9.3.1, GraphPad Software, San Diego, CA, USA). The differences of protein fermentation metabolites on cytokine production, cell viability, and ROS production were assessed by One-way analysis of variance (ANOVA) with Dunnett as post-test. Šídák’s multiple comparisons test was applied in Figures 7 and 8 to see the differences from media or LPS, respectively, with no added fatty acids. * p < 0.05, ** p < 0.01, *** p < 0.001, **** p < 0.0001.

## Funding and Acknowledgement

We thank Anni Mehlsen, Mads Tranholm Bruun, and Conan Christian Cassidy at the Section of Experimental Animal Models (University of Copenhagen, Denmark) for assisting with animal handling during the study. Zhuqing Xie is funded by China Scholarship Council (NO. 202006150033) for the PhD study at the University of Copenhagen.

## List of abbreviations

AHR: aryl hydrocarbon receptor
B: butyrate
BMDCs: bone marrow-derived dendritic cells
BCFAs: branched-fatty acids
DCs: dendritic cells
Dusp1: Dual Specificity Phosphatase 1
GM-CSF: granulocyte-macrophage colony stimulating factor
GPR: G-protein coupled receptors
HDACs: histone deacetylase
H_2_S: hydrogen sulfide
IB: iso-butyrate
IDO1: indoleamine 2,3-dioxygenase 1
IN: indole
IV: iso-valerate
MB: 2-methylbutyrate
MFI: mean fluorescent intensity
N: ammonium chloride
pC: p-Cresol
PH: phenol
pre-B: pre-treated with butyrate
pre-V: pre-treated with valerate
ROS: reactive oxygen species
SCFAs: short-chain fatty acids
S: sodium hydrosulfide
TLR: Toll-like receptors
V: valerate
7-AAD: 7-Aminoactinomycin D

## Author contribution

ZQX, DSN, and HF formulated the idea; ZQX and HF designed this study; ZQX carried out most of the experiments and analyzed the data; DBE and PRJ helped and performed parts of some experiments; ZQX and HF wrote the manuscript; ZQX, PRJ, DBE, DSN, and HF discussed and interpreted data; All authors critically revised and approved the final version of the manuscript.

## Supplementary

**Supplementary figure 1:**
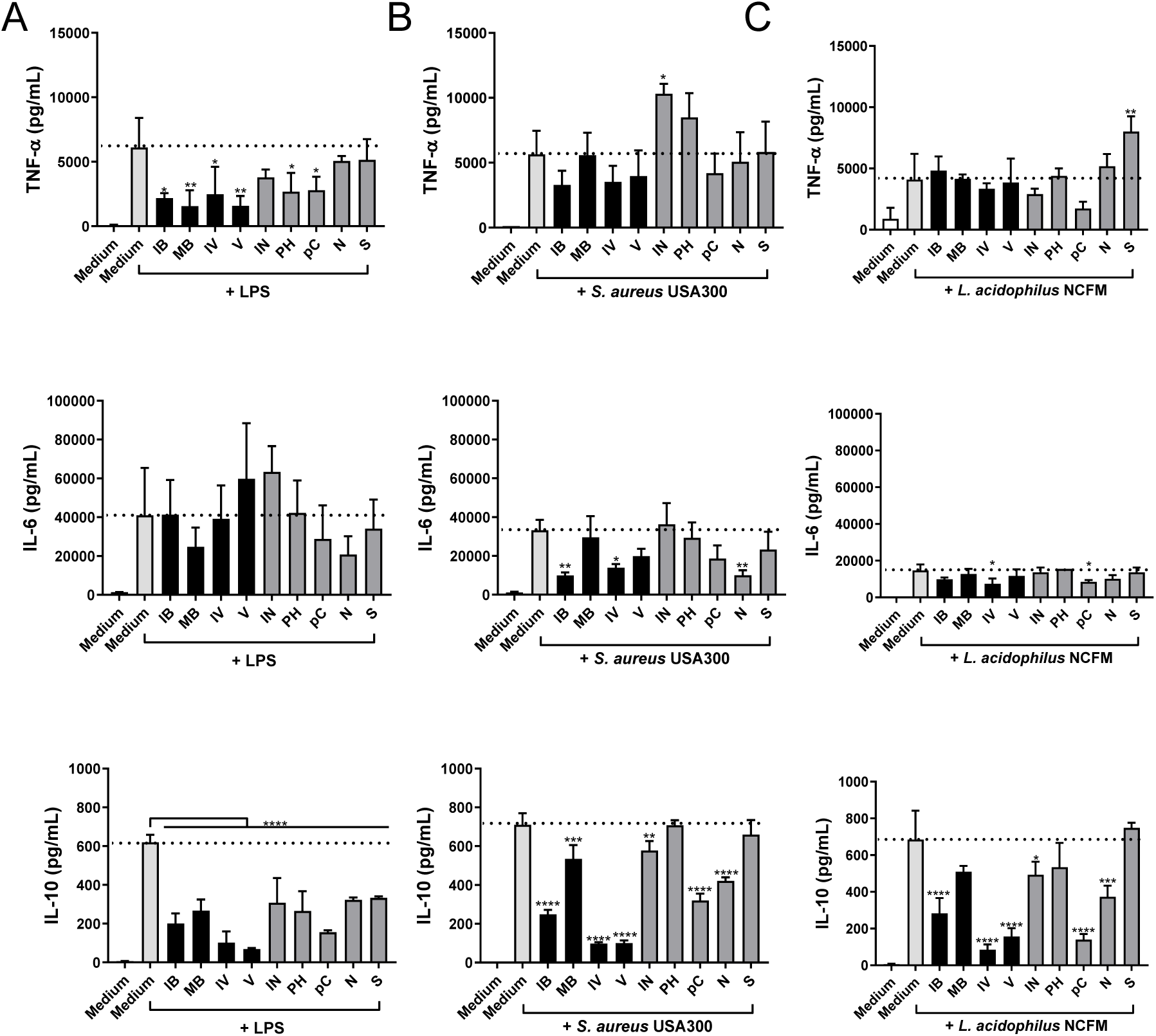
The effect of protein fermentation products on LPS **(A)**, *S. aureus* USA300 **(B)**, and *L. acidophilus* NCFM **(C)** induced TNF-α, IL-6, and IL-10 production. The compounds (1mM) were added to BMDC 30 min prior to the bacterial stimuli. After 20 hr incubation supernatants were harvested and cytokine concentration was determined by ELISA after stimulation. Differences in the production of cytokine were assessed by one-way analysis of variance (ANOVA) followed by Dunnett (* p < 0.05, ** p < 0.01, *** p < 0.001, **** p < 0.0001). IB, isobutyrate; MB, 2-methylbutyrate; IV, isovalerate; V, valerate; IN, indole; PH, phenol; pC, p-Cresol; N, ammonium chloride; S, sodium hydrosulfide.

**Supplementary figure 2:**
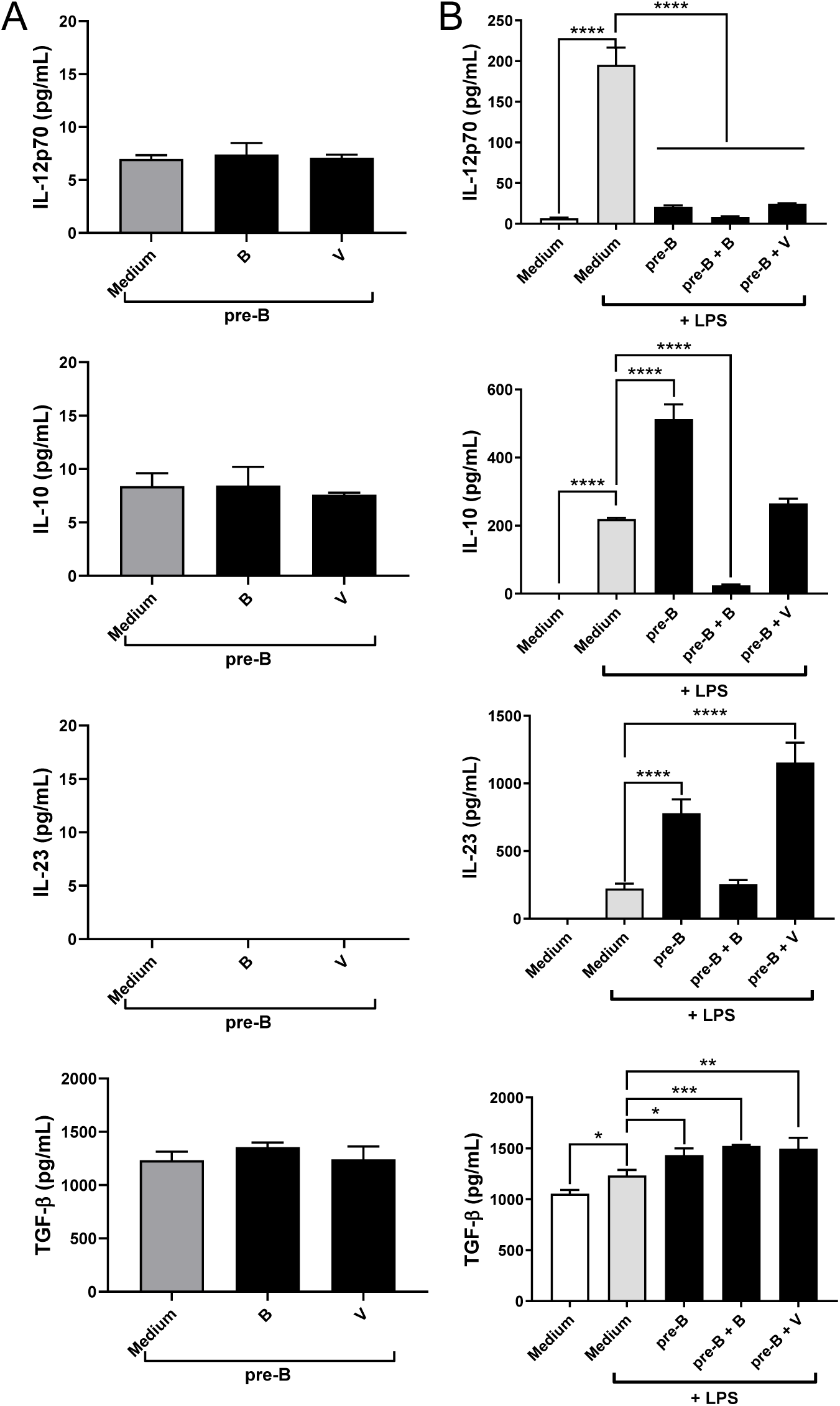
The production of cytokines in BMDCs pre-treated with butyrate (pre-B) for 2 days prior to harvest of cells and when further added butyrate (B) or valerate (V) 30 min before stimulation with medium **(A)** or LPS **(B)**. Cells were incubated for 20 h and supernatants were harvested and analyzed for cytokine concentrations. Statistics: Dunnett’s multiple comparisons test. Asterisks show differences between LPS-stimulated cells with no added fatty acids. * p < 0.05, ** p < 0.01, *** p < 0.001, **** p < 0.0001.

